# KAI2 regulates seedling development by mediating light-induced remodelling of auxin transport

**DOI:** 10.1101/2021.05.06.443001

**Authors:** Maxime Hamon-Josse, Jose Antonio Villaecija-Aguilar, Karin Ljung, Ottoline Leyser, Caroline Gutjahr, Tom Bennett

## Abstract

The photomorphogenic remodelling of seedling growth upon exposure to light is a key developmental transition in the plant life cycle. The α/β-hydrolase signalling protein KARRIKIN-INSENSITIVE2 (KAI2), a close homologue of the strigolactone receptor DWARF14 (D14), is involved in this process, and *kai2* mutants have strongly altered seedling growth as a result^1^. KAI2 and D14 both act through the MAX2 (MORE AXILLARY BRANCHING2) F-box protein to target proteins of the SMAX1-LIKE (SUPPRESSOR OF MAX2 1) (SMXL) family for degradation, but the signalling events downstream of this step are unclear in both pathways^2^. Here, we show that *kai2* phenotypes arise because of a failure to downregulate auxin transport from the seedling shoot apex towards the root system, rather than a failure to respond to light *per se*. We demonstrate that KAI2 controls the light-induced remodelling of the PIN-mediated auxin transport system in seedlings, promoting the reduction of PIN3, PIN4, and PIN7 abundance in older tissues, and the increase of PIN1, PIN2, PIN3, and PIN7 abundance in the root meristem, consistent with transition from elongation-mediated growth in the dark to meristematically-mediated growth in the light. We show that removing PIN3, PIN4 and PIN7 from *kai2* mutants, or pharmacological inhibition of auxin transport and synthesis, is sufficient to suppress most *kai2* seedling phenotypes. KAI2 is not required for the light-mediated changes in PIN gene expression but is required for the changes in PIN protein abundance at the plasma membrane; we thus propose that KAI2 acts to promote vesicle trafficking, consistent with previous suggestions about D14-mediated signalling in the shoot^3^.

## RESULTS AND DISCUSSION

### KAI2 mediates light induced remodelling of seedling development

The roles of KAI2 in the correct photomorphogenic development of hypocotyls and cotyledons in the light have previously been described^3,4^, as have the roles of KAI2 in the regulation of root development^5^. However, it has not been clear how these phenotypes are related to each other. Since *kai2* mutants display stronger phenotypes in younger seedlings, particularly in the roots^5^, we hypothesized that *kai2* phenotypes arise from sluggish adaption to the light, rather than a long-term inability to grow correctly in the light. To test this idea, we grew wild-type (Col-0) and *kai2-2* seedlings in the dark for 4 days (4dd), before tracking their development 2 days (4dd/2dl) and 4 days (4dd/4dl) after the transition to the light. In wild-type seedlings, the transfer to the light causes a rapid cessation of hypocotyl growth by 4dd/2dl, followed by a slight increase in hypocotyl growth and the development of both adventitious and lateral roots by 4dd/4dl (Figure 1, B, C). Consistent with previous studies, we found that *kai2* mutants have a wild-type hypocotyl phenotype in the dark but fail to stop growing after transition to light between 4dd and 4dd/2dl (Figure 1A). Similarly, we observed that *kai2* mutants initiate more adventitious roots than wild-type between 4dd and 4dd/2dl (Figure 1B). The increased adventitious roots in *max2* mutants, which block both KAI2 and strigolactone (SL) signalling^1^ were previously reported to be the result of decreased SL signalling^6^, although a more recent report has suggested that KAI2 signalling is more important for this phenotype^7^. Our results are consistent with this, and we also observed increased junction roots in light-grown *kai2* relative to wild-type, rather than in SL mutants (Supplemental Figure 1A). We also observed that lateral root formation is greater in *kai2* than wild-type between 4dd and 4dd/2dl, but sustained outgrowth does not differ significantly (Figure 1B). This phenotype arises because *kai2* mutants initiate more lateral root primordia that show increased emergence in comparison to wild-type (Supplemental Figure 1B). Growth in light conditions also promotes an increase in root hair density and root hair length in wild-type compared to dark-grown seedlings; but we observed that *kai2* mutants fail to develop root hairs normally, as previously reported^5^, independently of the light status (Supplemental Figure 1C, D). Thus, the *kai2* phenotype is not simply a failure to adapt to the light; *kai2* mutants over-respond to light exposure in the initiation of new roots. We also grew the *smax1-2 smxl2-1* (*s1s2*) mutant lacking the proteolytic targets of KAI2 activity^8^. The *s1s2* mutant phenotypes are similarly complex; *s1s2* hypocotyls seem to over-respond to light exposure, but they do not initiate adventitious roots in the light, while the number of lateral roots initially increases strongly, but fails to show a sustained increase (Figure 1A, B, C). Overall, the role of KAI2 signalling in photomorphogenesis seems to be more concerned with the correct spatial patterning of growth responses, rather than the response to light *per se*.

**Figure 1:**
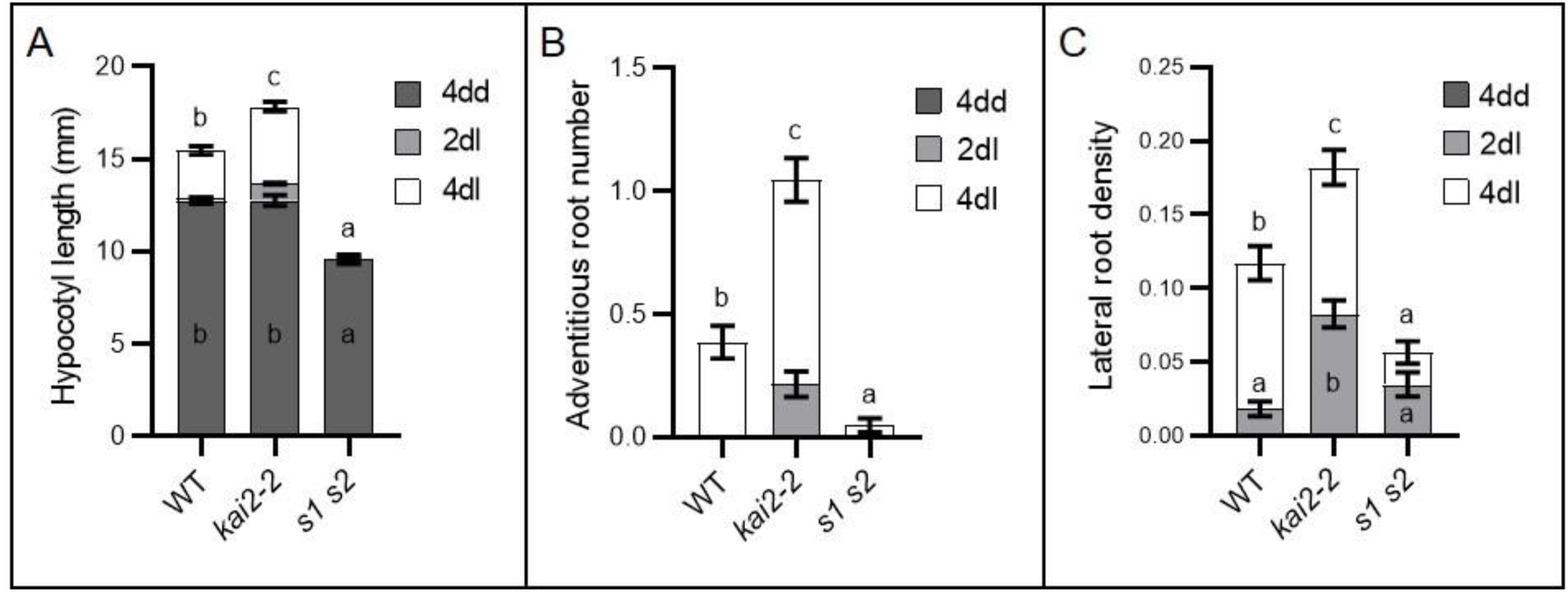
KAI2 mediates light induced remodelling of seedling development. (A-C) Hypocotyl length (A), adventitious root number (B), and lateral root density (C) in wild-type (WT), *kai2-2*, and *smax1-2 smxl2-1* (*s1 s2*) seedlings after 4 days growth in the dark (4dd), and after subsequent transfer to normal light conditions for 2 and 4 days (2dl and 4dl). Three independent experimental replicates with comparable results were performed (n=43-91 per experiment). Statistical groups were determined by one-way ANOVA with post hoc Tukey HSD (CI 95%). Error bars represent ± s.e.m.

### KAI2 modulates auxin distribution in the seedling

Given the prominent role of auxin in hypocotyl elongation and root growth^9,10^, and its known roles downstream of light perception^11,12^, we hypothesized that the *kai2* phenotype might arise due to perturbations in auxin response. To test this idea, we examined expression of the *DR5v2:GFP* auxin reporter in the relevant tissues after transfer to light. We found that auxin response is increased in the hypocotyls, adventitious root primordia and emerged lateral roots and primordia in *kai2*, consistent with the idea that auxin response is perturbed in these mutants (Figure 2A-D; Supplemental Figure 2A, B). We tested whether this altered response might be caused by increased auxin synthesis in *kai2* mutants, by directly measuring auxin levels. In seedlings dark-grown for 4 days, auxin levels in *kai2* and wild-type are identical (Figure 2E), but in 4dd/1dl seedlings, auxin levels in *kai2* seedlings were reproducibly higher than in wild-type. These data could therefore be consistent with increased auxin synthesis causing the *kai2* phenotype. Interestingly, however, this increase was larger in the hypocotyl/root compartment compared to the cotyledons (Figure 2F). It is also notable that in the roots of 5-day old light-grown (5dl) seedlings, auxin levels were similarly increased in *kai2* mutants relative wild-type, but so were levels in the *d14-1* SL receptor mutant, which does not have the same root phenotypes as *kai2-2* (Figure 2G)^5^. Conversely, the *s1s2 (smax1-2 smxl2-1)* and *smxl6-4 smxl7-3 smxl8-1 (s678)* triple mutant (which lacks the proteolytic targets of D14 activity) have the same auxin levels as wild-type, but have dramatic root phenotypes not present in wild-type^5^. Thus, changes in auxin synthesis alone are unable to explain the specific phenotypes observed in *kai2* and *smax1 smxl2* mutants.

**Figure 2:**
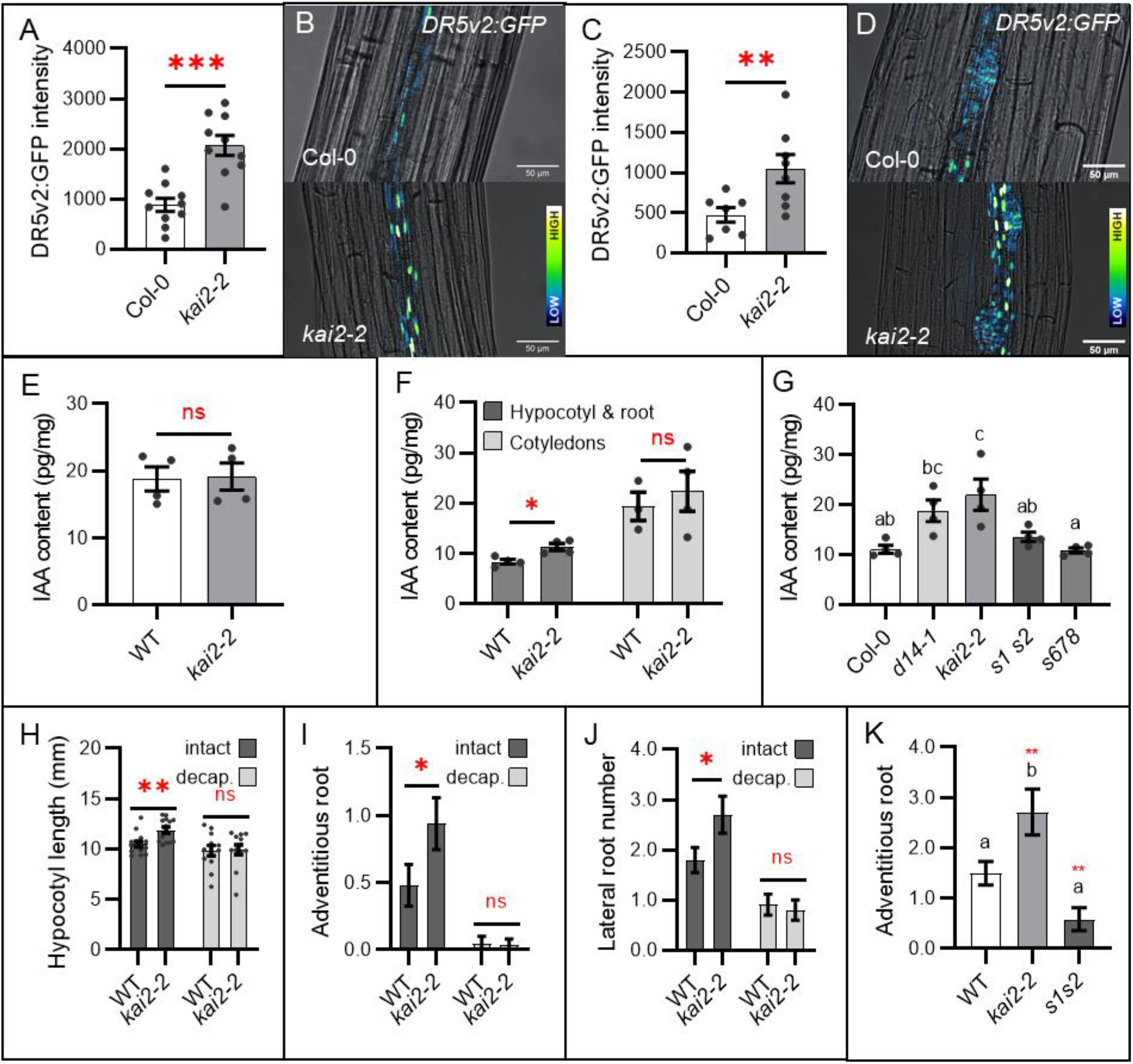
KAI2 modulates auxin distribution in the seedling. (A-D) Auxin response (*DR5v2:GFP* fluorescence intensity) in seedlings grown 4 days in the dark followed by 4 days of normal light-growth, in the hypocotyl (A, B) and adventitious root primordia (C, D). (A) and (C) show GFP quantification in the hypocotyl. For (A) n=12-15 seedling per genotype; for (C) n= 7-9 primordia from 5 different seedlings for each genotype; Two independent experimental replicates with comparable results were performed. (*,**,***) p-value ≤ (0.05, 0.01, 0.001) indicates differences compared to wild-type (Welch’s t-test). Error bars represent ± s.e.m. (B) and (D) show representative microscopy images with bright field (grey) and GFP signals represented with false colour with dark blue as low signal intensity and bright white as high signal intensity. Scale bars represent 50 μm. (E-G) IAA quantification (pg IAA per mg of tissue, pg/mg) in whole seedlings grown for 4 days in the dark (E), or roots grown 5 days under normal light conditions (G), or in cotyledons and hypocotyl/root sections of seedlings grown for 4 days in the dark and subsequently transferred to normal light conditions for 1 day (F). (n=3-4 pools of 30 seedlings). Statistical groups indicated by letters were determined by one-way ANOVA with post hoc Tukey HSD (CI 95 (*,**,***) p-value ≤ (0.05, 0.01, 0.001) indicates differences compared to wild-type (Welch’s t-test). Error bars represent ± s.e.m. (H-J) Measurement of (H) hypocotyl, (I) adventitious root number, and (J) lateral root number of seedlings grown for 4 days in the dark and subsequently undergoing apex decapitation (light grey) or left intact (dark grey) prior to transfer to normal light conditions for 3 days. Data are from two independent experimental replicates with comparable results (n=12-14 seedlings per genotype and treatment for each experiment). (*,**,***) p-value ≤ (0.05, 0.01, 0.001) indicates differences compared to wild-type (Welch’s t-test). Error bars represent ± s.e.m. (K) Adventitious root development in 11 days old de-rooted seedlings from three independent experimental replicates with comparable results (n = 17-21 per experiment). (*,**,***) p-value ≤ (0.05, 0.01, 0.001) indicates differences compared to wild-type (Welch’s t-test). Error bars represent ± s.e.m.

Given the altered partitioning of auxin in *kai2* mutants at 4dd/1dl (Figure 2F), we reasoned that the effects of KAI2 on seedling development might relate more to altered auxin distribution than to simply increased auxin levels. Specifically, we hypothesized that failure to downregulate auxin transport from the cotyledons towards the root might account for all *kai2* seedling phenotypes. To test this idea, we first used microsurgical approaches. We de-capitated wild-type and *kai2-2* seedlings at 4dd by removing the seedling apex, and then assessed their phenotype at 4dd/3dl. This treatment was sufficient to restore the *kai2* hypocotyl, adventitious root, and lateral root phenotypes to wild-type (Figure 2H-J; Supplemental Figure 2D-E), consistent with apically-derived auxin driving these effects. We next de-rooted etiolated Col-0, *kai2-2*, and *s1s2* seedlings after 4dd and 3dl and assessed their adventitious root phenotype at 4dd/7dl. This treatment amplified the *kai2* phenotype but had no effect on the *s1s2* phenotype (Figure 2K; Supplemental Figure 2C). These data are thus consistent with increased auxin transport and accumulation in the hypocotyls of *kai2*, and reduced auxin transport and accumulation in *s1s2*. Thus, the phenotype of KAI2 appears to be associated with increased auxin synthesis and increased rootward auxin flux, similar to the shoot branching phenotypes of strigolactone mutants^3,13,14^.

### KAI2 regulates remodelling of auxin transport at the dark-light transition

We therefore hypothesized that the altered auxin transport in *kai2* might be caused by increased abundance of members of the PIN family of auxin efflux carriers, which play major roles in mediating directional auxin transport^15^, and have been implicated in the phenotypic effects of strigolactone signalling^3,13,14,16^. We examined the abundance of PIN proteins in wild-type hypocotyls in the dark (4dd) and after transition to the light (4dd 3dl) using GFP protein fusions. PIN3, PIN4 and PIN7 are all highly abundant at 4dd, but greatly reduced by 4dd/3dl (Figure 3A). PIN1 and PIN2 are absent from the hypocotyl under both conditions (Supplemental Figure 3A). We also examined PIN protein abundance in roots. PIN2 accumulation at the plasma membrane has previously been reported to be induced by light^17^, and we found that PIN1 plasma membrane abundance, and indeed total PIN1 abundance also increased in the root meristem after exposure to light (Figure 3D, E). PIN4 was difficult to detect, but PIN3 and PIN7 are highly abundant throughout the length of 4dd roots. In older root tissues, their abundance declines by 4dd/1dl, but in the root meristem and elongation zone their abundance is maintained or increases (Supplemental Figure 3B-D). Thus, after the dark-light transition, there is a major re-organization of the seedling auxin transport network. Expression of the PIN genes, as assessed by qRT-PCR, reflects these changes in PIN abundance, with *PIN3, PIN4* and *PIN7* all downregulated and *PIN1* upregulated at 4dd/1dl and 4dd/3dl relative to 4dd (Figure 3F; Supplemental Figure 3H).

**Figure 3:**
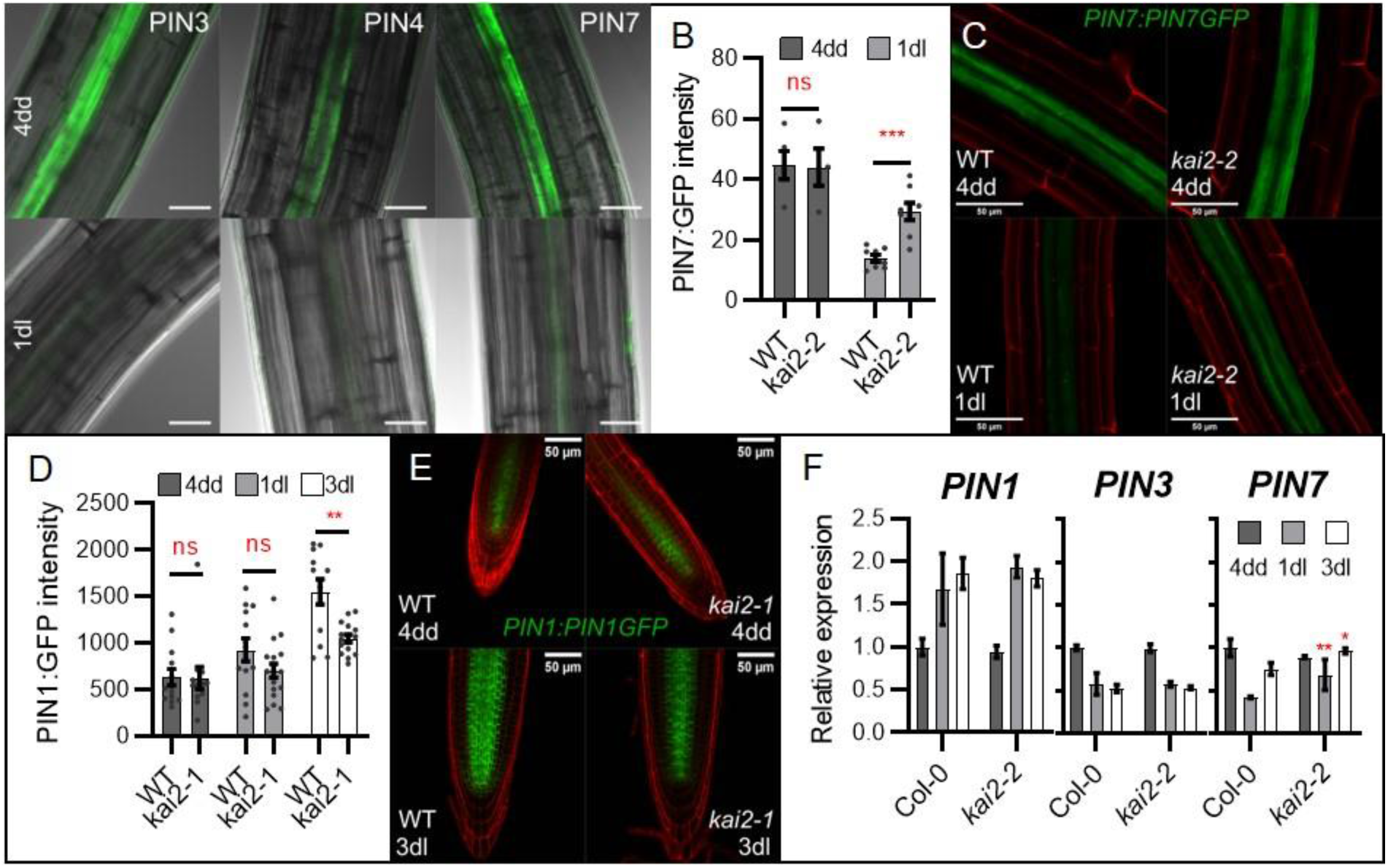
KAI2 regulates non-transcriptional re-modelling of auxin transport at the dark-light transition. (A) PIN3-GFP, PIN4-GFP, and PIN7-GFP abundance in hypocotyls of wild-type seedlings after 4 days growth in the dark (4dd, top row), and subsequent transfer to normal light conditions for 3 days (3dl, bottom row). Images overlay bright field (grey) and GFP signals (green). Scale bars represent 30 μm. (B-C) PIN7-GFP abundance quantification (B) and representative images (C) in the old differentiation zone of wild-type or *kai2* roots after 4 days growth in the dark (4dd), and subsequent transfer to normal light conditions for 1 day (1dl). (n=4-8 seedling per genotype per experiment); two independent experimental replicates with comparable results were performed. (*, **,***) p-value ≤ (0.05, 0.01, 0.001) indicates differences compared to wild-type (Welch’s t-test). Error bars represent ± s.e.m. (B) Microscopy images overlay propidium iodide (red) and GFP signals (green). Scale bars represent 50 μm. (D-E) PIN1:GFP abundance quantification (D) and representative images (E) in root meristem zone of wild-type or *kai2* seedlings after 4 days growth in the dark (4dd), and subsequent transfer to normal light conditions for 1 and 3 days (1dl, 3dl). (n=12-17 seedling per genotype per experiment); two independent experimental replicates with comparable results were performed. (*,**,***) p-value ≤ (0.05, 0.01, 0.001) indicates differences compared to wild-type (Welch’s t-test). Error bars represent ± s.e.m. (E) Microscopy images overlay propidium iodide (red) and GFP signals (green). Scale bars represent 50 μm. (F) Expression of *PIN* genes relative to the reference gene *UBC10* in wild-type and *kai2-2* seedlings after 4 days growth in the dark (4dd), and subsequent transfer to normal light conditions for 1 and 3 days (1dl, 3dl). For each gene, expression is normalised to the expression in wild type at 4dd). (n=3 biological samples collected by pooling ~16 seedlings per genotype and time-point); (*,**,***) p-value ≤ (0.05, 0.01, 0.001) indicates differences compared to wild-type (Welch’s t-test). Error bars represent ± s.e.m.

We next tested whether this re-organization is delayed in *kai2* mutants, consistent with the changes in auxin distribution we observed (Figure 2). In hypocotyls we observed a delay in the reduction of PIN3 abundance in some experiments, but this was not consistent, and the imaging was not straightforward. We therefore focused on PIN abundance in the root. We observed no difference in PIN7 abundance in mature root tissues between wild-type and *kai2-2* at 4dd, but there was a clear failure to decrease PIN7 abundance in *kai2-2* at 4dd/1dl relative to wild-type (Figure 3B, C). Conversely, for PIN1 abundance in the meristem zone, we observed the opposite; there was no difference between wild-type and *kai2-1* at 4dd, but there was delay in the increase of PIN1 abundance at 4dd/1dl and 4dd/3dl (Figure 3D, E). These differences are long-lasting, and increased PIN7 abundance and decreased PIN1 abundance can be detected in young light-grown seedlings (Supplemental Figure 3E-G). Thus, *kai2* mutants show a general reduction in the rate that the auxin transport system is re-modelled after transition to the light. However, we did not observe any clear differences in PIN gene expression in *kai2* relative to Col-0 at 4dd, 4dd/1dl or 4dd/3dl (Figure 3F; Supplemental Figure 3H). This suggests that KAI2 regulates PIN abundance non-transcriptionally, perhaps through effects on the cycling of PIN proteins to and from the plasma membrane. This may be consistent with the recent data showing that MAX2 and *rac*-GR24 signalling inhibits the inhibitory effect of auxin on PIN endocytosis^16^. Although these effects were suggested to reflect the output of strigolactone signalling, they might equally reflect outputs of KAI2 signalling, given the prominent use of *max2* mutants and *rac*-GR24 in these experiments^1,2^. Either way, this idea certainly parallels the previously described role of D14-mediated strigolactone signalling in the regulation of PIN endocytosis and auxin transport in the shoot system^3^, and suggests that this function is conserved between the D14 and KAI2 signalling pathways.

### The phenotypic effects of KAI2 signalling are mediated by PIN-mediated auxin transport

Our data support the idea that the altered auxin distribution we observed in *kai2* (Figure 2) is caused by a failure to remodel the auxin transport system after dark-light transition (Figure 3), and strongly suggest that this causes the accompanying failure to remodel seedling development after dark-light transition. To test this model, we used the auxin transport inhibitor 1-N-naphthylphthalamic acid (NPA)^18^ to try and rescue the *kai2* phenotype. Consistent with our model, treatments in the range of 0.1-1μM NPA were sufficient to reduce the light grown hypocotyl, adventitious root, and lateral root phenotypes of *kai2* to a wild-type level (Figure 4A, B; Supplemental Figure 4A). We were able to achieve the same effect using the auxin synthesis inhibitor L-kynurenine^19^ at a concentration of 10nM (Supplemental Figure 4B, C). To provide independent verification of these results, we crossed *kai2-2* to the *pin3-3 pin4-3 pin7-1* triple mutant^20^. The verified quadruple mutant (*k2 pin347*) restored the light grown hypocotyl phenotype to a wild-type level (Figure 4C). The adventitious root phenotype of *kai2* is also rescued in the quadruple mutant (Figure 4D), but the lateral root phenotype is harder to assess since *pin3 pin4 pin7* drastically reduced lateral root formation in a wild-type background.

**Figure 4:**
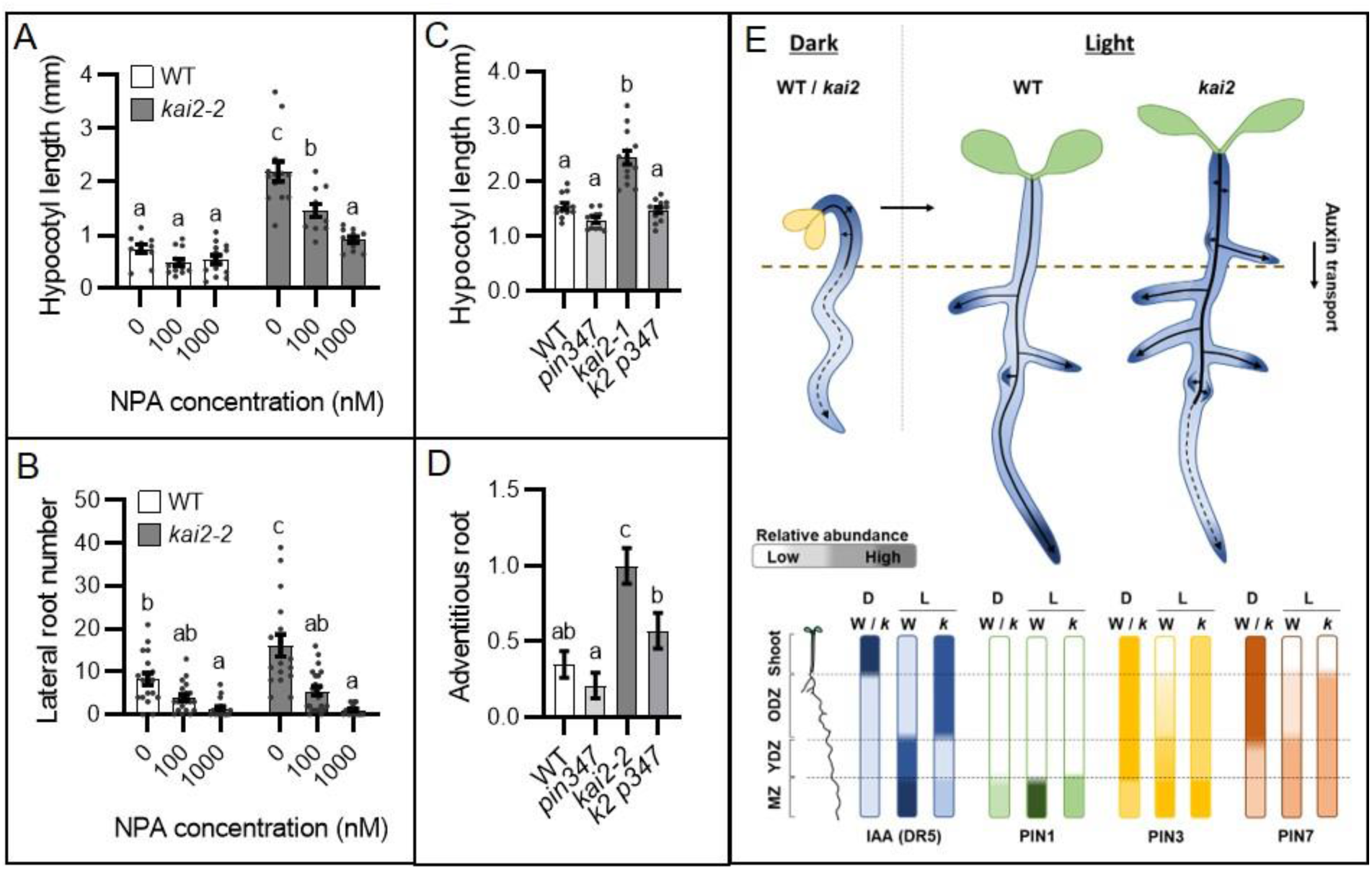
The phenotypic effects of KAI2 signalling are mediated by PIN-mediated auxin transport. (A-B) Effect of auxin transport inhibitor 1-N-naphthylphthalamic acid (NPA) on hypocotyl length (A) and lateral root number (B) in 10 day-old wild-type and *kai2-2* seedlings. For (A) n=11-13; for (B) n= 18-22 seedlings per treatment and per genotype). Experiments were independently repeated three and two times respectively with comparable results. Statistical groups indicated by letters were determined by one-way ANOVA with post hoc Tukey HSD (CI 95%). Error bars represent ± s.e.m. (C-D) Hypocotyl length (C) and adventitious root number (D) in 10 day-old wild-type, *kai2-2, pin3-3 pin4-3 pin7-1 (pin347) and kai2-2 pin3-3 pin4-3 pin7-1 (k2 p347)* seedlings. For (C) n=11-14; for (D) n= 40-45 seedlings per genotype). Experiments were independently repeated two times with comparable results. Statistical groups indicated by letters were determined by one-way ANOVA with post hoc Tukey HSD (CI 95%). Error bars represent ± s.e.m. (E) Proposed model for light-induced remodelling of the auxin transport system to regulate seedling development. During skotomorphogenesis, seedlings have a strong rootward auxin transport from the shoot apex, mediated by high PIN3 and PIN7 abundance, which drives the elongation growth of the hypocotyl and primary root. At the transition to photomorphogenic development, KAI2 mediates the rapid remodelling of the PIN-mediated auxin transport system, with a reduction of PIN3 and PIN7 abundance in older tissues, and increased PIN1, PIN3 and PIN7 abundance in the root meristem, to promote a meristematically-mediated growth in the light. In *kai2* mutants, the failure to remodel the auxin transport system transition leads to excess auxin in the shoot and older root tissues promoting continued hypocotyl elongation and increased adventitious and lateral roots growth in the shootward part of the root, with reduced auxin delivery to the primary root meristem.

Taken together, these data are consistent with our model that KAI2 mediates light-induced remodelling of the auxin transport system to regulate seedling development. In the dark, there is a strong auxin transport connection between the shoot apex and cotyledons and the root, with high PIN3, PIN4 and PIN7 abundance along the main shoot-root axis of the plant, and low PIN1/PIN2 abundance in the root meristem (Figure 4E). Intriguingly, this auxin transport system does not seem to be particularly important for skotomorphogenic development (consistent with^9^), since *pin3 pin4 pin7* etiolates normally. Its function might relate more to the delivery of auxin to the root system for future growth, than to growth in the dark (Figure 4E). After the transition to light, this system is rapidly remodeled, with turnover of PIN3, PIN4 and PIN7 in the hypocotyl and older root tissues, and upregulation of PIN1, PIN2, PIN3, PIN4 and PIN7 in the root meristem (Figure 4F), PIN1 upregulation may act to ‘inject’ auxin into the meristem from the rest of the root, driving cell division, consistent with the strong increase in meristem size that occurs in the light^21^. Meanwhile PIN2, PIN3, PIN4 and PIN7 upregulation, in concert with AUX1 (Villaécija-Aguilar et al., accompanying manuscript), promotes the ‘reflux’ of auxin from the root cap to the epidermis, which drives elongation zone activity and root hair development (Figure 4E). Collectively, this remodelling ‘kick-starts’ the meristematic ‘engine’ of growth, allowing a move away from primarily elongation-driven growth in the dark. However, in *kai2* mutants, the failure to quickly re-model the system leads to continued auxin transport from cotyledons into the hypocotyl, delaying expansion of the cotyledons, and promoting continued hypocotyl elongation, and the formation of adventitious roots (Figure 4E). This excess auxin reaches the roots, where it promotes increased lateral root initiation and emergence. Interestingly, we found that overall transport of auxin from the shoot-to-root junction to the tip of *kai2* roots is decreased relative to Col-0 (Supplemental Figure 4D), presumably because it is diverted into the lateral roots, which show increased auxin reporter activity relative to wild-type (Supplemental 2A, B). Consistent with this reduced overall transport, we found that auxin reporter activity is *diminished* in the primary root meristem of *kai2-2*, the opposite to that seen in the lateral roots (Supplemental Figure 4F, G). This reduced auxin delivery to the primary root meristem likely accounts for some of the phenotypes observed in the primary root in *kai2*^5^.

### New perspectives on KAI2 signalling

The data presented here provide a holistic explanation for the role of KAI2 in seedling growth. It has long been speculated that KAI2 targets the SMAX1 (SUPPRESSOR OF MAX2 LIKE 1) and SMXL2 proteins for degradation, analogous to the D14-SMXL7/D53 interaction^2^, and recent data provide confirmation of this idea^8,22^. However, apart from an increase in ethylene biosynthesis in the root^23,24^, KAI2-mediated signalling events downstream of SMAX1/SMXL2 remain unclear. There are some well-established genes upregulated in response to KAI2 signalling, including its homologue *DWARF14-LIKE2* (*DLK2*) and *KAI2-UPREGULATED F-BOX1* (*KUF1*)^1^, but the function of these genes remains enigmatic, and DLK2 has no obvious role in development^25^. Our data suggest that analogous to the role of D14 in the shoot^3,26^, KAI2 regulates the cycling of PIN auxin transporters between the endosomal system and the plasma membrane. By promoting this efficient cellular re-modelling of the auxin transport system, KAI2 promotes a larger-scale re-modelling of the auxin transport system, allowing seedlings to undergo rapid changes in growth in response to light exposure. In the shoot, the mechanism by which D14-mediated signalling events in the nucleus lead to changes in vesicle trafficking is unclear^27^. It is clear that the effects of both D14/KAI2 on PIN proteins do not involve changes in PIN gene expression, and are unaffected by cycloheximide^3^, but it is possible that D14/KAI2 may rather upregulate some aspect of vesicle trafficking instead. There is certainly increasing evidence for the role of SMXL7/D53 as regulators of transcription^28^, but no clear connection to vesicle trafficking has been identified. However, the ability to use a seedling-based system will greatly simplify future investigations of this mechanism.

## MATERIALS AND METHODS

### Plant Materials

*Arabidopsis thaliana* genotypes were in Columbia-0 (Col-0) or Landsberg *erecta* (Ler) parental backgrounds. The *max2-1*^29^, *max2-8*^30^, *max4-5*^13^, *d14-1*^1^, *kai2-1*^1^ *kai2-2* [backcrossed 6x to Col-0]^14^, *smax1-2 smxl2-1^31^, smxl6-4 smxl7-3 smxl8-1^32^, DR5v2:GFP^33^, pin3-3 pin4-3 pin7-1^20^, PIN1:PIN1-GFP^34^, PIN3:PIN3-GFP, PIN4:PIN4-GFP, PIN7:PIN7-GFP^35^, PIN2:PIN2-GFP^36^*, lines have all been previously described.

New genotypes were assembled by crossing relevant existing genotypes and required homozygous lines were identified using visible, fluorescent, or selectable markers or using PCR genotyping.

### Growth conditions

Seedlings for phenotypic analysis, dissections, pharmacological treatments, auxin quantification, qPCR and confocal imaging were grown in axenic culture. Seeds were surface sterilized using 2 hours vapor sterilization method (3 ml of HCl 37% in 100 ml bleach), then sown onto 0.8% agar-solidified *Arabidopsis thaliana* salts (ATS) media (pH 5.6)^37^ with 1% (w/v) sucrose, in square petri dishes (12 x 12cm, 60 ml media per plate), and stratified in the dark at 4°C for 2-3 days.

For plants grown in normal light conditions, plates were oriented vertically, and seedlings grown for 6-10 days in growth chamber under a 16-h/8-h light/dark cycle (20°C/18°C) with light provided by fluorescent tubes (120 μmolm^-2^s^-1^). In experiments with transfer from dark to light conditions, after stratification, plates were placed for 8 hours at 120 μmolm^-2^s^-1^ light / 20°C to promote germination and then placed in complete darkness for 4 days at 20°C in a black plastic box in a growth chamber. After 4 days in darkness plates were transferred in normal light conditions described above (16h/8h light/dark cycle (20°C/18°C) / 120 μmolm^-2^s^-1^), for an additional 1 to 6 days of growth.

### Phenotypic analysis

Measurements of seedlings were made at various time points described in the text. A dissecting microscope was used to count emerged adventitious and lateral roots in each root system. Plants were then imaged using a flatbed scanner, and primary root and hypocotyl length was quantified from the resulting images using Fiji (https://imagej.net/Fiji/Downloads). Lateral root density was quantified as the number of lateral roots per mm of primary root. Lateral root and Adventitious root primordia numbers were scored by observing DR5v2:GFP as a primordia marker in Col-0 and *kai2*-2 with a Laser-scanning confocal microscope LSM880 upright (see laser microscopy section below). For analysis of root hair length, *Arabidopsis thaliana* seeds were grown in axenic conditions on 12×12cm square Petri dishes containing 60 ml in ½ Murashige & Skoog medium, pH 5.8 (Duchefa), supplemented with 1% sucrose and solidified with 1.5% agar. Prior to sawing seeds were surface sterilized by washing with 1 ml of 70% (v/v) ethanol and 0.05% (v/v) Triton X-100 with gentle mixing by inversion for 6 minutes at room temperature, followed by one wash with 96% ethanol and 3 washes with sterile distilled water. Plants were stratified at 4°C for 3 days in the dark, and then transferred to a growth cabinet at 22°C and placed vertically under light provided by fluorescent tubes (120 μmolm^-2^s^-1^) for 3 hours to promote germination. Subsequently they were grown in the same growth chamber for 5 days in complete darkness in a black plastic box, in light (16-h/8-h light/dark cycle), or in dark/light (16-h/8-h light/dark cycle) with roots covered with a black paper to keep them in dark, while the aerial part remained illuminated.

Root images were taken at 5 days post germination with a Zeiss Stereo Discovery V8 microscope (Carl Zeiss, Germany) equipped with a Zeiss Axiocam 503 color camera (Carl Zeiss, Germany). The number of root hairs was determined by counting the root hairs between 2 and 3 mm from the root tip on each root, and root hair length was measured for 10 root hairs per root in a minimum of 8 roots per genotype and condition using Fiji (https://imagej.net/Fiji/Downloads) according to Villaécija-Aguilar et al (2021)^38^.

### Seedling dissections

Four-days-old etiolated seedlings were dissected in-situ on agar plates using a very sharp scalpel. Decapitation assays were performed by removing the seedling meristem and cotyledons at the junction of the cotyledons and hypocotyl. De-rooting assays were performed by excising the root at the root-shoot junction.

### Pharmacological Experiments

For pharmacological experiments, 1000X stock solutions 1-N-naphthylphthalamic acid (NPA) (Duchefa), and L-kynurenine (Sigma-Aldrich), by dissolving the appropriate mass of the compound in a 2% DMSO, 70% ethanol solution. From these stocks 60μl/plate was added to hand-warm ATS-agar media prior to pouring plates. Control plates contained 60μl/plate of 2% DMSO, 70% ethanol solvent control solution. Seed were either germinated directly on plates containing to the pharmacological treatments, or were transferred after initial growth on plain plates, as indicated in the text.

### Free IAA Determination

For the first experiment (Figure 2F), seedlings were grown for 4 days on ATS-agar medium with sucrose in the dark, and then transferred to the light. Some seedlings were dissected immediately, with the cotyledons separated from the hypocotyl + roots, and flash frozen in liquid nitrogen. The other seedlings were dissected after 1 further day of growth in the light, and then dissected in the same way. There were 4 biological samples for each genotype and time point, each containing pooled tissue from 60 seedlings. For the second experiment (Figure 2G) seedlings were grown for 6 days on ATS-agar medium with sucrose in the light, at which point the roots were dissected from the seedlings, and flash frozen. There were 4 biological samples for each genotype and time point, each containing pooled tissue from 60 seedlings. From these samples (10-20 mg fresh weight) IAA was purified and analyzed by gas chromatography-tandem mass spectrometry (GC-MS/MS) as described in Andersen et al (2008)^39^ with minor modifications. To each sample, 500 pg ^13^C_6_-IAA was added as an internal standard before extraction.

### Auxin transport assay

For the auxin transport assay, *Arabidopsis thaliana* seed sterilization and seedling growth were performed as described for root hair measurements in light conditions. An agar droplet containing 100 nM ^3^H-IAA (Hartmann analytic) and solvent (DMSO) or 10 μM NPA (Olchemim) in DMSO was applied below the aligned root-shoot junctions of 5 days post germination Arabidopsis seedlings. 18 hours after the treatment, the amount of radioactivity was quantified in a 5 mm apical segment as previously described in Lewis and Muday (2009)^40^.

### Laser Scanning Confocal Microscopy

To visualize fluorescent reporter lines Laser-scanning confocal microscopy was performed on either Zeiss LSM700 or LSM880 imaging system with a 20X lens. Tissues were stained with propidium iodide (10ug/ml) and mounted on glass slides. GFP excitation was performed using a 488 nm laser, and fluorescence was detected between 488 and 555nm. Propidium iodide excitation was performed using a 561 nm laser, and fluorescence was detected between above 610nm. The same detection settings were used for all images captured in a single experiment. GFP quantification was performed on non-saturated images, using Zeiss ‘ZEN’ software.

For the different GFP lines (DR5v2:GFP, PIN1-GFP, PIN2-GFP, PIN3-GFP, PIN4-GFP, and PIN7-GFP) fluorescence was quantified in regions of interest either in the hypocotyl, the shoot-root junction, the older differentiated zone (ODZ, between the first two emerged lateral roots), the middle differentiated zone (MDZ) between the last emerged LR and the first LR primordia), the young differentiated zone (YDZ, in the root hair elongation zone), or in the meristem zone including columella and quiescent centre nuclei as appropriate.

### RNA extraction and gene expression analysis

For expression analysis of PIN genes, Col-0 and *kai2-2* seedlings were grown for 4 days on ATS-agar medium with sucrose in the dark, and then transferred to the light for 0, 1 or 3 additional days of growth. For each time point and genotype, 3 biological samples were collected by pooling ~16 seedlings, which were then flash-frozen in liquid nitrogen. Total RNA was extracted using RNeasy Plant Mini kits (Qiagen), and then DNase treated using Turbo DNA-free kit (Ambion), both as per manufacturer’s instructions. RNA was quantified using a Nanodrop 1000. For cDNA synthesis, Superscript (Invitrogen) II was used to reverse transcribe 500 ng of total RNA according to manufacturer’s instructions. Quantification of transcript levels was carried out using SYBR Green reactions with 5 ng cDNA in a 20 μl volume on a Light Cycler 480 II (Roche) relative to the reference gene *UBC10* (*POLYUBIQUITIN10*, At4g05320). Three technical replicates were run for each biological replicate and averaged. Calculation of the expression levels was done using the ΔΔCt method^41^. Primers used were:

*PIN1*-F: 5′-CAGTCTTGGGTTGTTCATGGC-3′; *PIN1*-R: 5′-ATCTCATAGCCGCCGCAAAA-3′. *PIN3*-F: 5′-CCATGGCGGTTAGGTTCCTT-3′; *PIN3*-R: 5′-ATGCGGCCTGAACTATAGCG-3′. *PIN4*-F: 5′-AATGCTAGAGGTGGTGGTGATG-3′; *PIN4*-R: 5′-TAGCTCCGCCGTGGAATTAG-3′. *PIN7*-F: 5′-GGTGAAAACAAAGCTGGTCCG-3′; *PIN7*-R: 5′-CCGAAGCTTGTGTAGTCCGT-3′ UBQ10-F: 5’-GGTTTGTGTTTTGGGGCCTTG-3’; UBQ10-R: 5’-CGAAGCGATGATAAAGAAGAAGTTCG-3’

### Statistical Analyses

Statistical analyses were performed in R-studio, using one-way Analysis of Variance (ANOVA), followed by Tukey HSD post hoc test (CI 95%) or Welch’s t-test, with (*,**,***) p-value ≤ (0.05, 0.01, 0.001) indicating differences between genotype or conditions, as appropriate. The test(s) performed for each figure is indicated in legend.

## Acknowledgments

TB is supported by BBSRC (BB/R00398X/1) and CG by the Emmy Noether Program of the Deutsche Forschungsgemeinschaft (GU1423/1-1). OL is supported by Gatsby Charitable Foundation grant GAT3272C. KL is supported by grants from the Swedish Research Council, The Knut and Alice Wallenberg Foundation, and the Swedish Governmental Agency for Innovation Systems. KL thanks Roger Granbom for excellent technical assistance.

## Author contributions

TB, OL and CG designed the study. MHJ, JAVA, KL and TB planned and carried out experiments and analyzed data. MHJ and TB wrote the manuscript with input from all authors.

## FIGURE LEGENDS

**Supplemental Figure 1:**
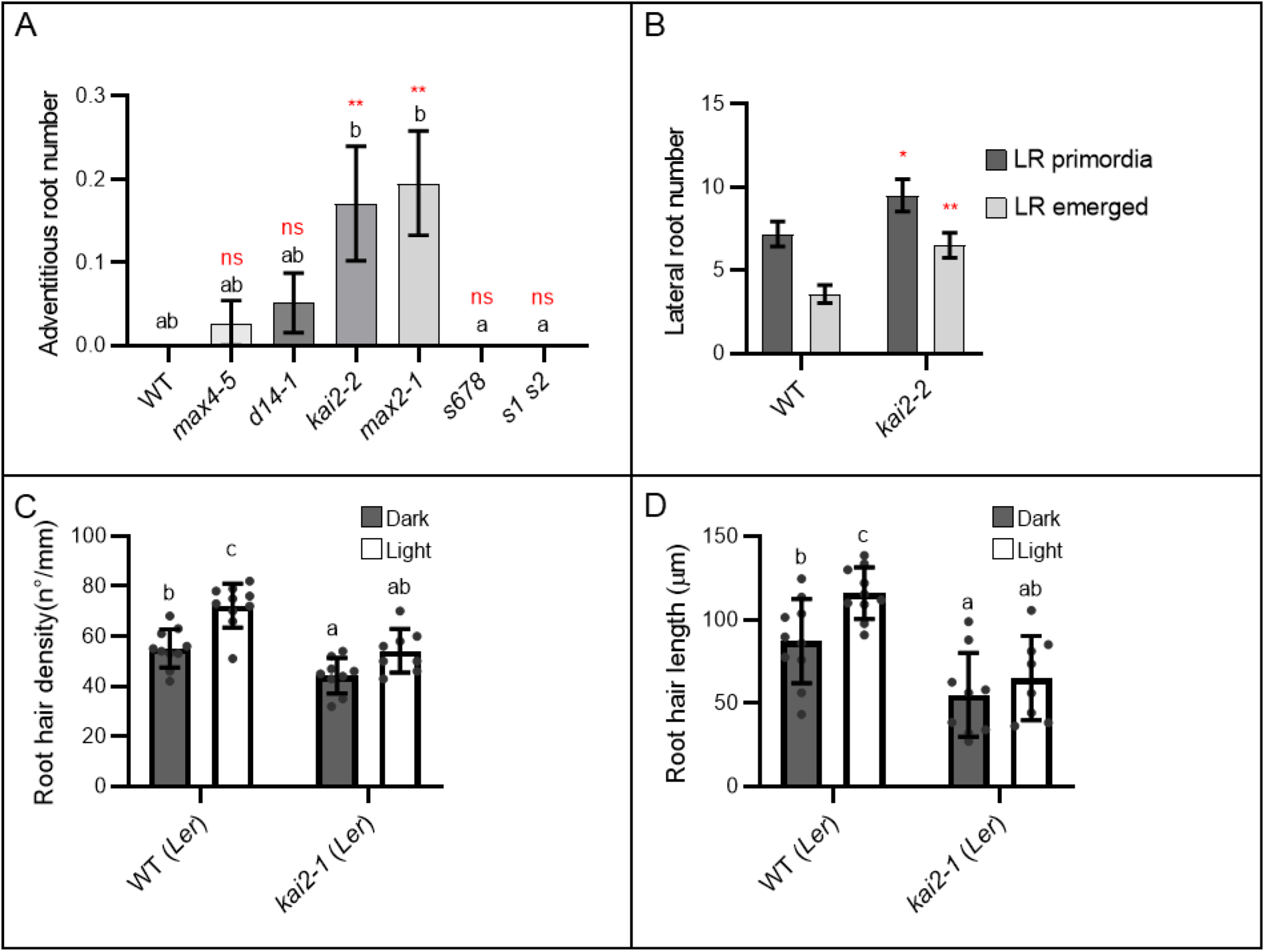
KAI2 mediates light induced remodelling of seedling development. (A) Adventitious and junction root number in 10 days old light-grown seedlings of SL and KL signalling mutants. Data are from four independent experimental replicates with comparable results (n =19-44 seedlings per experiment). Statistical groups were determined by one-way ANOVA with post hoc Tukey HSD (CI 95%); (*,**,***) p-value ≤ (0.05, 0.01, 0.001) indicates differences compared to wild-type (Welch’s t-test). Error bars represent ± s.e.m. (B) Number of lateral root primordia and emerged lateral roots in 10 day old light-grown seedlings from three independent experimental replicates with comparable results (n =11-30 seedlings per experiment); (*,**,***) p-value ≤ (0.05, 0.01, 0.001) indicates differences compared to wild-type (Welch’s t-test). Error bars represent ± s.e.m. (C-D) Root hair density (C) and length (D) in 5 day-old seedlings grown in the dark (dark), or in normal light conditions (light). Data are from 2 independent experimental replicates with comparable results (n =8-10 seedlings per experiment for each condition). Statistical groups were determined by one-way ANOVA with post hoc Tukey HSD (CI 95%).

**Supplemental Figure 2:**
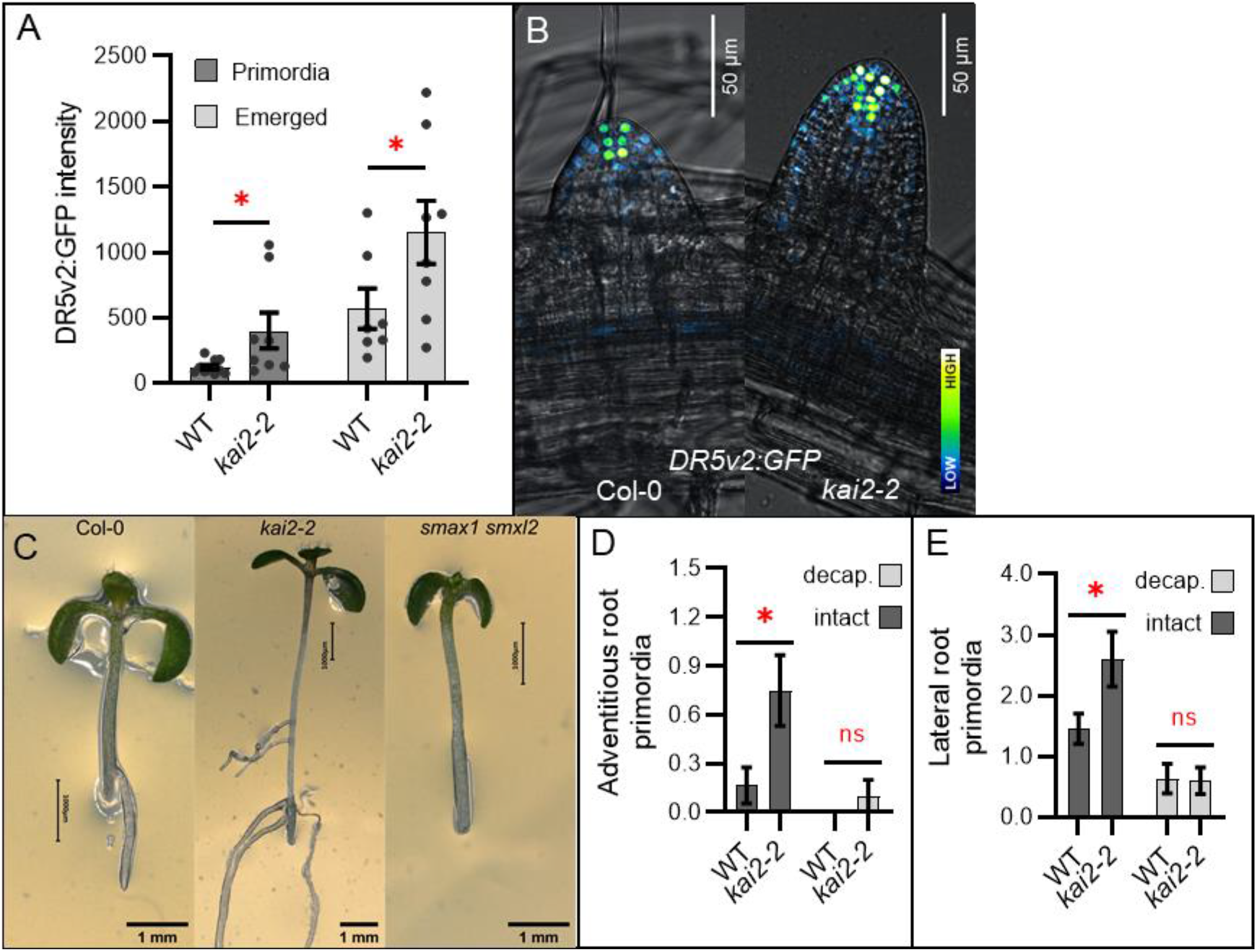
KAI2 modulates auxin distribution in the seedling. (A-B) Auxin response quantification (*DR5v2:GFP* fluorescence intensity) (A) and representative images (B) in lateral roots of WT and *kai2-2* seedlings grown for 6 days under normal light conditions. (n=8-10 lateral roots from 5 different seedlings for each genotype). Three independent experimental replicates with comparable results were performed. (*,**,***) p-value ≤ (0.05, 0.01, 0.001) indicates differences compared to wild-type (Welch’s t-test). Error bars represent ± s.e.m. (B) Microscopy images overlay bright field (grey) and GFP-derived signals. GFP signal is represented with false-colour with dark blue as low signal intensity and bright white as high signal intensity. Scale bars represent 50 μm. (C) Images showing adventitious root outgrowth 11 days post-germination in de-rooted and etiolated seedlings dark-grown for 4 days then transferred to normal light conditions for 3 days before de-rooting. Scale bars represent 1 mm. (D-E) Adventitious (D) and lateral (E) root primordia number in seedlings grown in the dark for 4 days and subsequently undergoing apex de-capitation (light grey) or left intact (dark grey) before transfer to normal light conditions for 3 days. Data are from two independent experimental replicates with comparable results (n=12-14 seedlings per genotype and treatment for each experiment). (*,**,***) p-value ≤ (0.05, 0.01, 0.001) indicates differences compared to wild-type (Welch’s t-test). Error bars represent ± s.e.m.

**Supplemental Figure 3:**
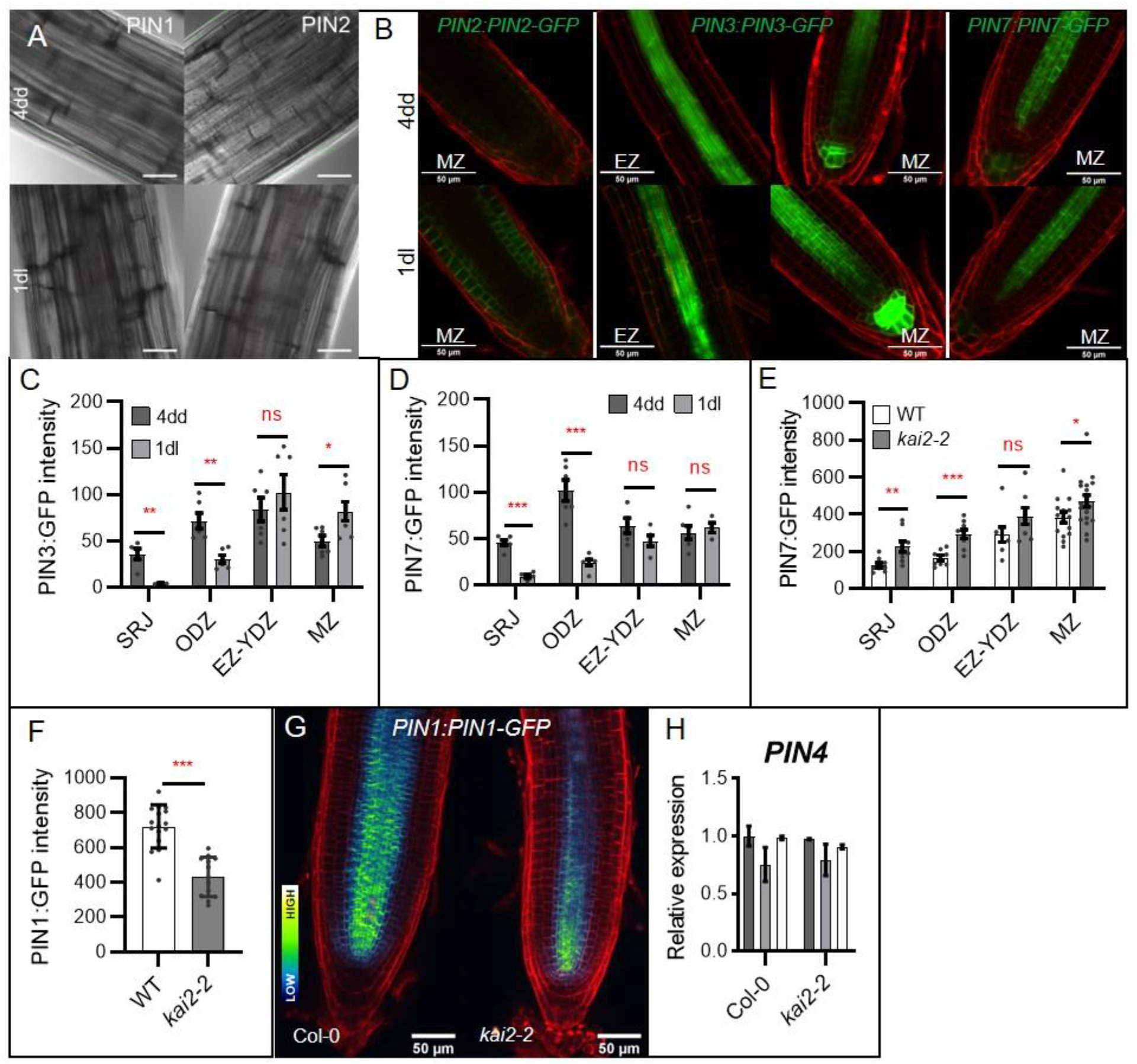
KAI2 regulates re-modelling of auxin transport at the dark-light transition. (A) PIN1-GFP and PIN2-GFP abundance in hypocotyl tissues of wild-type seedlings after 4 days growth in the dark (4dd, top row), and subsequent transfer to normal light conditions for 3 days (3dl, bottom row). Images overlay bright field (grey) and GFP signals (green). Scale bars represent 30 μm. (B) PIN2-GFP, PIN3-GFP and PIN7-GFP abundance in meristem zone (MZ) and elongation zone (EZ) of wild-type seedlings after 4 days growth in the dark (4dd, top row), and subsequent transfer to normal light conditions for 1 day (1dl). Microscopy images overlay propidium iodide (red) and GFP signals (Green). Scale bars represent 50 μm. (C-D) Quantification of PIN3-GFP (C) and PIN7-GFP (D) signals in the shoot-root junction (SRJ), older differentiation zone (ODZ), junction of the elongation and young differentiation zone (EZ-YDZ), and meristem zone (MZ) of wild-type seedlings after 4 days growth in the dark (4dd), and subsequent transfer to normal light conditions for 1 day (1dl). (n=4-7 per tissue and experiment); two independent experimental replicates with comparable results were performed. (*,**,***) p-value ≤ (0.05, 0.01, 0.001) indicates differences compared to wild-type (Welch’s t-test). Error bars represent ± s.e.m. (E) Quantification of PIN7-GFP signal in the shoot-root junction (SRJ), older differentiation zone (ODZ), junction of the elongation and young differentiation zone (EZ-YDZ), and meristem zone (MZ) of 6 day-old wild-type and *kai2-2* seedlings grown under normal light conditions (n=8-18 per tissue, per genotype, and per experiment); two independent experimental replicates with similar results were performed. (*,**,***) p-value ≤ (0.05, 0.01, 0.001) indicates differences compared to wild-type (Welch’s t-test). Error bars represent ± s.e.m. (F-G) PIN1-GFP quantification (F) and representative microscopy images (G) in the meristem zone of 6 day-old wild-type and *kai2-2* seedlings grown under normal light conditions (n=13-16 per genotype per experiment); three independent experimental replicates with comparable results were performed. (*,**,***) p-value ≤ (0.05, 0.01, 0.001) indicates differences compared to wild-type (Welch’s t-test). Error bars represent ± s.e.m. Microscopy images overlay propidium iodide staining (red) and GFP-derived signal represented with false colour with dark blue as low signal intensity and bright white as high signal intensity. Scale bars represent 50 μm. (H) Expression of *PIN4* gene relative to the reference gene *UBC10* in wild-type and *kai2-2* seedlings after 4 days growth in the dark (4dd), and subsequent transfer to normal light conditions for 1 and 3 days (1dl, 3dl). Expression is normalised to the expression in wild type at 4dd. (n= 3 biological samples collected by pooling ~16 seedlings per genotype and time-point). (*,**,***) p-value ≤ (0.05, 0.01, 0.001) indicates differences compared to wild-type (Welch’s t-test). Error bars represent ± s.e.m.

**Supplemental Figure 4:**
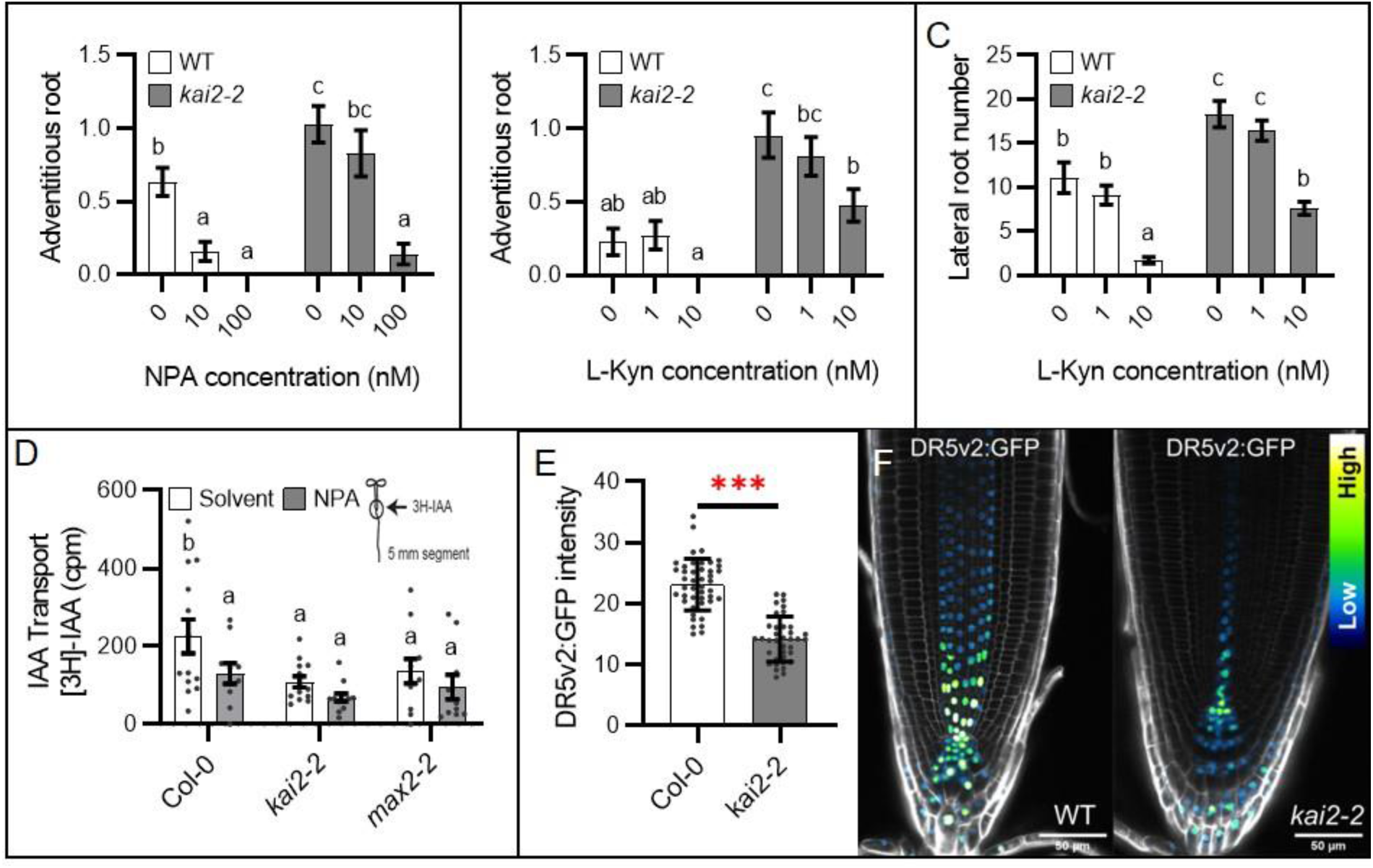
The phenotypic effects of KAI2 signalling are mediated by PIN-mediated auxin transport. (A) Effect of auxin transport inhibitor 1-N-naphthylphthalamic acid (NPA) on adventitious and junction root number in 10 day-old wild-type and *kai2-2* seedlings (n=38-42 seedlings per treatment and per genotype). Experiment was independently repeated two times with comparable results. Statistical groups indicated by letters were determined by one-way ANOVA with post hoc Tukey HSD (CI 95%). Error bars represent ± s.e.m. (B-C) Effect of auxin biosynthesis inhibitor L-kynurenine (L-KYN) on adventitious (B) and lateral root (C) number in 10 day-old wild-type and *kai2-2* seedlings (n=21-22 seedlings per treatment and per genotype). Experiment was independently repeated three times with comparable results. Statistical groups indicated by letters were determined by one-way ANOVA with post hoc Tukey HSD (CI 95%). Error bars represent ± s.e.m. (D) Basipetal ^3^H-IAA auxin transport in 5 day-old seedlings. Arrow indicates the site of ^3^H-IAA or solvent application at the shoot-root junction, radioactivity was measured in a 5 mm segment at the primary root meristem. (n=10-13 seedlings per treatment and genotype) Experiment was independently repeated two times with comparable results. Statistical groups indicated by letters were determined by one-way ANOVA with post hoc Tukey HSD (CI 95%). Error bars represent ± s.e.m. (E-F) Auxin response (*DR5v2:GFP* fluorescence intensity) in root meristem zone of seedlings in normal light-grown conditions. (E) Shows GFP signal quantification and (F) representative images. (n=37-44 seedling per experiments). Two independent experimental replicates with comparable results were performed. (*,**,***) p-value ≤ (0.05, 0.01, 0.001) indicates differences compared to wild-type (Welch’s t-test). Error bars represent ± s.e.m. Microscopy images overlay propidium iodide staining (grey) and GFP-derived signal represented with false colour with dark blue as low signal intensity and bright white as high signal intensity. Scale bars represent 50 μm.

